# Optical activation of TrkA signaling

**DOI:** 10.1101/287409

**Authors:** Liting Duan, Jen M. Hope, Shunling Guo, Qunxiang Ong, Amaury François, Luke Kaplan, Grégory Scherrer, Bianxiao Cui

## Abstract

Nerve growth factor/tropomyosin receptor kinase A (NGF/TrkA) signaling plays a key role in neuronal development, function, survival, and growth. The pathway is implicated in neurodegenerative disorders including Alzheimer’s disease, chronic pain, inflammation, and cancer. NGF binds the extracellular domain of TrkA, leading to the activation of the receptor’s intracellular kinase domain. TrkA signaling is highly dynamic, thus mechanistic studies would benefit from a tool with high spatial and temporal resolution. Here we present the design and evaluation of four strategies for light-inducible activation of TrkA in the absence of NGF. Our strategies involve the light-sensitive protein *Arabidopsis* cryptochrome 2 (CRY2) and its binding partner CIB1. We demonstrate successful recapitulation of native NGF/TrkA functions by optical induction of plasma membrane recruitment and homo-interaction of the intracellular domain of TrkA. This approach activates PI3K/AKT and Raf/ERK signaling pathways, promotes neurite growth in PC12 cells, and supports the survival of dorsal root ganglion neurons in the absence of NGF. This ability to activate TrkA using light bestows high spatial and temporal resolution for investigating NGF/TrkA signaling.

---

During embryonic development, nerve growth factor (NGF) plays a critical role in supporting neuronal differentiation, survival, and plasticity ^1^. In adults, NGF supports neural maintenance and repair, and deficits in NGF signaling have been implicated in several neurodegenerative disorders including Alzheimer’s and Parkinson’s diseases ^2–4^. Aberrantly elevated activity of NGF is also involved in chronic inflammatory and neuropathic pain disorders ^5,6^. The tropomyosin receptor kinase A (TrkA) has been shown to be the primary receptor of NGF in both neuroprotection and pain. Thus, therapeutic strategies that aim to augment NGF/TrkA signaling have been pursued for neurodegenerative diseases ^7–10^. For example, NGF gene therapies have been explored in clinical trials for Alzheimer’s disease ^11,12^, and small molecules agonists of TrkA are also being sought for potential use as therapies ^13,14^. On the other hand, therapeutic strategies that aim to block NGF/TrkA signaling are being explored as novel methods for pain management; Pfizer’s anti-NGF monoclonal antibody tanezumab^15–17^ received fast-track designation from the FDA for pain management in 2017 and has progressed to Phase 3 clinical trials.

However, the development of NGF-related drugs has been hindered partly by the incomplete understanding of NGF/TrkA signaling and its dynamics due to the lack of methods to precisely activate TrkA. It is widely accepted that NGF binds to the extracellular domain of TrkA, leading to its autophosphorylation and activation. There is less certainty regarding the identities of downstream signaling effectors or the composition of the signaling endosomes that transport TrkA after activation. TrkA signaling is dynamically regulated in space and time; the outcome of activation can differ based on subcellular environment, protein interaction partners, and the duration of activation. Thus, the ability to activate TrkA signaling with precise spatial and temporal control would be ideal for dissecting the discrete signaling events of NGF application.

The emergence of light-inducible tools presents new strategies to control intracellular activities with high spatial and temporal resolution. These optogenetic tools utilize photo-sensitive proteins that undergo light-gated protein-protein interactions, such as the light-oxygen-voltage (LOV) domain ^18^, phytochrome B ^19^, and cryptochrome 2 (CRY2) ^20^. Among them, *Arabidopsis* CRY2 can undergo both light-induced homo-oligomerization and hetero-dimerization with its binding partner CIB1 ^21^. Optogenetic strategies based on CRY2/CIB1 hetero-dimerization have been successfully developed to control a variety of intracellular activities, such as organelle transport ^22,23^, gene expression ^20,24^, inactivation or disruption of protein function ^21,25^, and activation of diverse intracellular kinases ^26–29^. The light-mediated homo-oligomerization of CRY2 has also been used to regulate intracellular signaling of kinases such as Raf ^30^ and transmembrane receptor kinases such as DCC ^31^. In particular, the use of CRY2 oligomerization-based optogenetic strategies to control the activation of TrkA has been explored in a previous work ^26^. This previously reported strategy appends CRY2 to the C-terminus of full length TrkA (flTrkA-CRY2), in which blue light induces flTrkA-CRY2 oligomerization to activate TrkA. The light-dependent activation of flTrkA-CRY2 was assayed by the nuclear translocation of extracellular signal-regulated kinase (ERK) but whether it fully recapitulates NGF functions is yet to be determined.

In this work, we design and compare four different strategies for light-induced TrkA activation. We find that light-induced homo-interaction of the intracellular domain of TrkA (iTrkA) is more effective than that of flTrkA in activating TrkA receptors and recapitulating NGF’s functions. Particularly, recruitment and homo-oligomerization of iTrkA on the cell membrane is most efficient in activating downstream signaling pathways, inducing neurite outgrowth in PC12 cells, and supporting the survival of DRG neurons in the absence of NGF. This new method of optical activation of TrkA will be a useful tool to investigate the dynamics of NGF/TrkA signaling and to facilitate therapeutic screening of small molecule drugs that target the intracellular domain of TrkA.

The TrkA receptor consists of three domains, the extracellular domain (ECD), a transmembrane domain (TM) and the intracellular domain (iTrkA) (Figure 1A). Dimeric NGF binds the extracellular domain of TrkA and induces receptor dimerization and conformational change. The catalytic intracellular domain is then autophosphorylated and initiates downstream signal transduction.^32,33^ Previous work has shown that chemical-induced dimerization of plasma membrane-targeted myr-iTrkA is sufficient to activate TrkA and results in differentiation in PC12 cells.^33^ Inspired by this work, we hypothesized that dimerization of the iTrkA domain using light-induced protein-protein interaction may achieve optical activation of TrkA and its downstream pathways. To build the light-controlled TrkA systems, we chose to use the CRY2-CIB1 dimerization and the CRY2-CRY2 homoligomerization systems.

**Figure 1:**
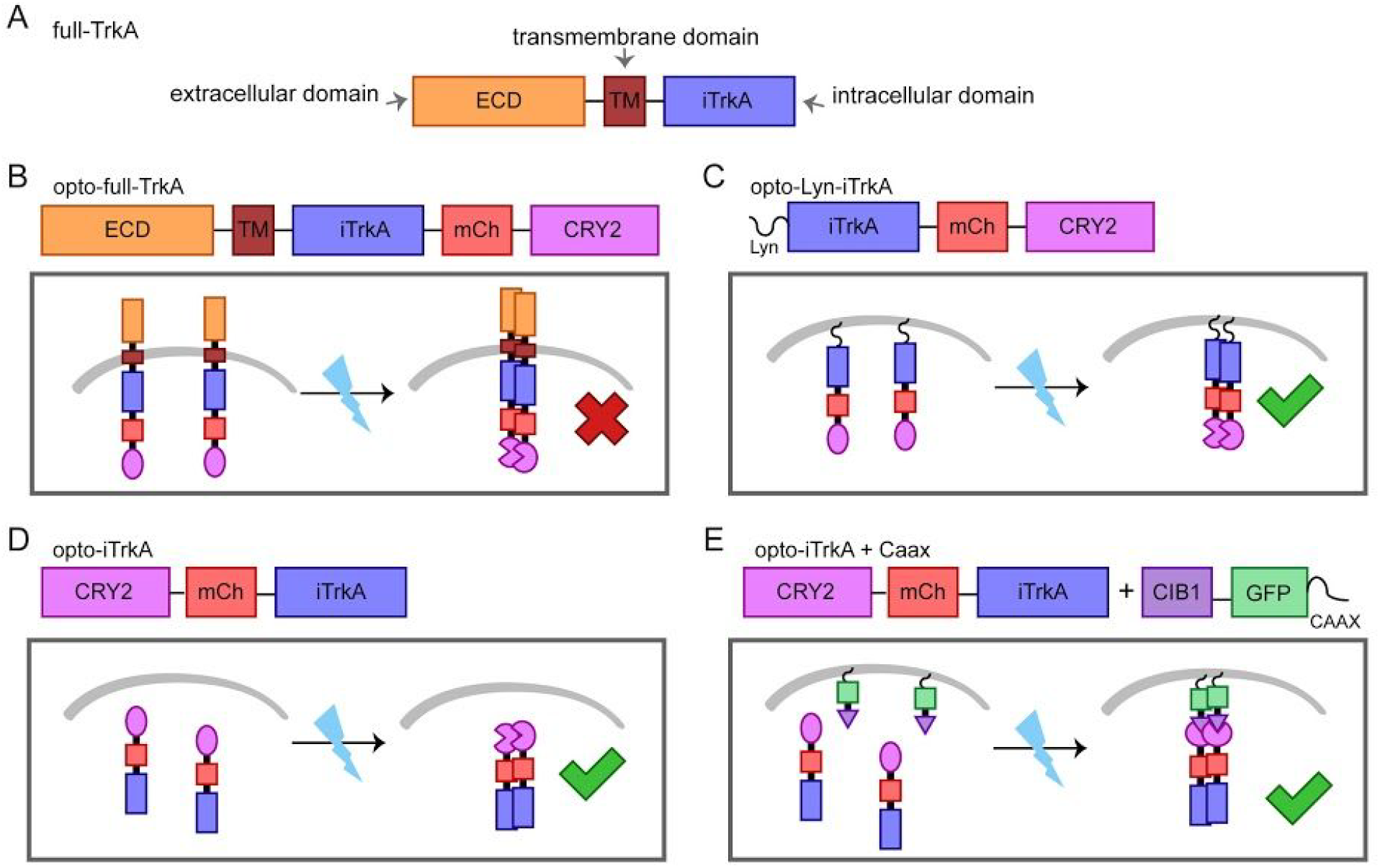
Design scheme of the light-controlled TrkA systems. We have constructed four methods of optogenetic activation of TrkA signaling by fusion to CRY2. The molecular architecture of each construct is detailed. (A) The full length TrkA receptor consists of an extracellular domain (ECD), a transmembrane domain (TM) and an intracellular domain (iTrkA). (B) Opto-full-TrkA is a fusion of CRY2 to the C-terminus of full length TrkA. (C) Opto-Lyn-iTrkA appends CRY2 to the C-terminal end of the intracellular domain of TrkA (iTrkA), and features an N-terminal Lyn membrane-targeting sequence. (D) Opto-iTrkA fuses CRY2 to the N-terminus of iTrkA. (E) opto-iTrkA+CAAX uses CIBN-CAAX, localized to the plasma membrane, in combination with opto-iTrkA to recruit iTrkA to the membrane in a light-dependent manner.

In order to achieve light-inducible TrkA activation, we designed four different strategies using CRY2. First, CRY2 was fused to the C-terminus of full-length TrkA to make opto-flTrkA. We expect that the fusion protein will be trafficked to the plasma membrane similar to endogenous TrkA, and that light-induced homo-oligomerization of CRY2 will initiate interaction between TrkA molecules (Figure 1B). This strategy has been reported previously ^26^. The second strategy fuses CRY2 to the C terminus of the truncated intracellular domain iTrkA, and is localized to the plasma membrane by an N-terminal Lyn tag (Lyn-iTrkA-CRY2, hereafter opto-Lyn-iTrkA). Light-induced CRY2 homo-interaction will induce oligomerization of iTrkA at the cell membrane (Figure 1C). Our third strategy uses a fusion of CRY2 to the N-terminus of iTrkA without any localization tag. We expect that blue light will induce the interaction of iTrkA in the cytosol (opto-iTrkA, Figure 1D). Finally, our last strategy uses co-expression of CRY2-iTrkA with CIB1-CAAX, which targets CIB1 to the plasma membrane via the CAAX motif ^34^. With this design, light stimulation induces both recruitment of CRY2-iTrkA to the cell membrane through CRY2-CIB1 dimerization and simultaneously induces iTrkA homo-interaction on the membrane through CRY2-CRY2 binding. We refer to the fourth strategy as opto-iTrkA+CAAX (Figure 1E). These four light-gated TrkA systems together are collectively denoted as optoTrkA systems henceforth.

In order to validate our optoTrkA systems, we examined signaling pathways downstream of TrkA. We first confirmed that our constructs could be expressed in cell culture, and that blue light stimulation induced membrane recruitment of iTrkA in cells expressing the opto-iTrkA+CAAX system (Figure S1). We next sought to assess whether blue light stimulation of optoTrkA systems would lead to the activation of pathways downstream of TrkA activation.

TrkA activation induces both the PI3K/AKT and the Raf/MEK/ERK signaling cascades. We assessed the activation of Raf/MEK/ERK by monitoring translocation of ERK2 from the cytosol to the nucleus, and probed the activation of PI3K/AKT by monitoring the translocation of the pleckstrin homology domain of AKT (PH_AKT1_) to the plasma membrane. NIH 3T3 cells were transfected with each of the four optoTrkA systems as well as ERK2-GFP or GFP-PH_AKT1_. All TrkA fusion proteins were tagged with mCherry (mCh). In the case of opto-iTrkA+CAAX, the fusion protein CIB1-CAAX was used, which contains no fluorescent reporter. mCherry fluorescence (excitation ~560 nm) was used to locate transfected cells under a microscope. Then 200 ms blue light pulses were delivered at 5 s intervals for up to 10 min to stimulate optoTrkA systems as well as to visualize the distribution of ERK2-GFP or GFP-PH_AKT1_.

In cells expressing opto-Lyn-iTrkA, cytosolic opto-iTrkA, or opto-iTrkA+CAAX, blue light stimulation induced visible recruitment of GFP-PH_AKT1_ to the plasma membrane (Fig 2A) and the nuclear translocation of ERK2-GFP (Fig 2B). On the other hand, opto-flTrkA did not induce visible changes in the spatial distribution of either GFP-PH_AKT1_ or ERK2-GFP. The translocation of GFP-PH_AKT1_ or ERK2-GFP was quantified to show the activation of PI3K/AKT or Raf/MEK/ERK respectively. Light-induced membrane recruitment of GFP-PH_AKT1_ was quantified as the ratio of average intensity at the membrane to the average cytosolic intensity after light stimulation, which was then normalized to the same ratio before light stimulation. Light-induced nuclear translocation of ERK2-GFP was measured similarly by the ratio of nuclear intensity to the average cytosolic intensity (see Methods for details). Quantitative analysis shows that blue light activation of opto-flTrkA did not induce a significant change in the membrane/cytosol and the nuclear/cytosol ratio for GFP-PH_AKT1_ and ERK2-GFP (Fig. 2C). In comparison, opto-Lyn-iTrkA, opto-iTrkA, and opto-iTrkA+CAAX, demonstrated significant blue-light induced increases in GFP-PH_AKT1_ and ERK2-GFP translocation. To confirm the pathway activation, we showed that the membrane translocation of GFP-PH_AKT1_ was inhibited by wortmannin, a PI3K inhibitor. Similarly, the nuclear translocation of ERK-GFP was blocked by adding Trametinib, a MEK inhibitor, in cells expressing opto-Lyn-iTrkA, opto-iTrkA or opto-iTrkA+CAAX (Figure S2).

**Figure 2:**
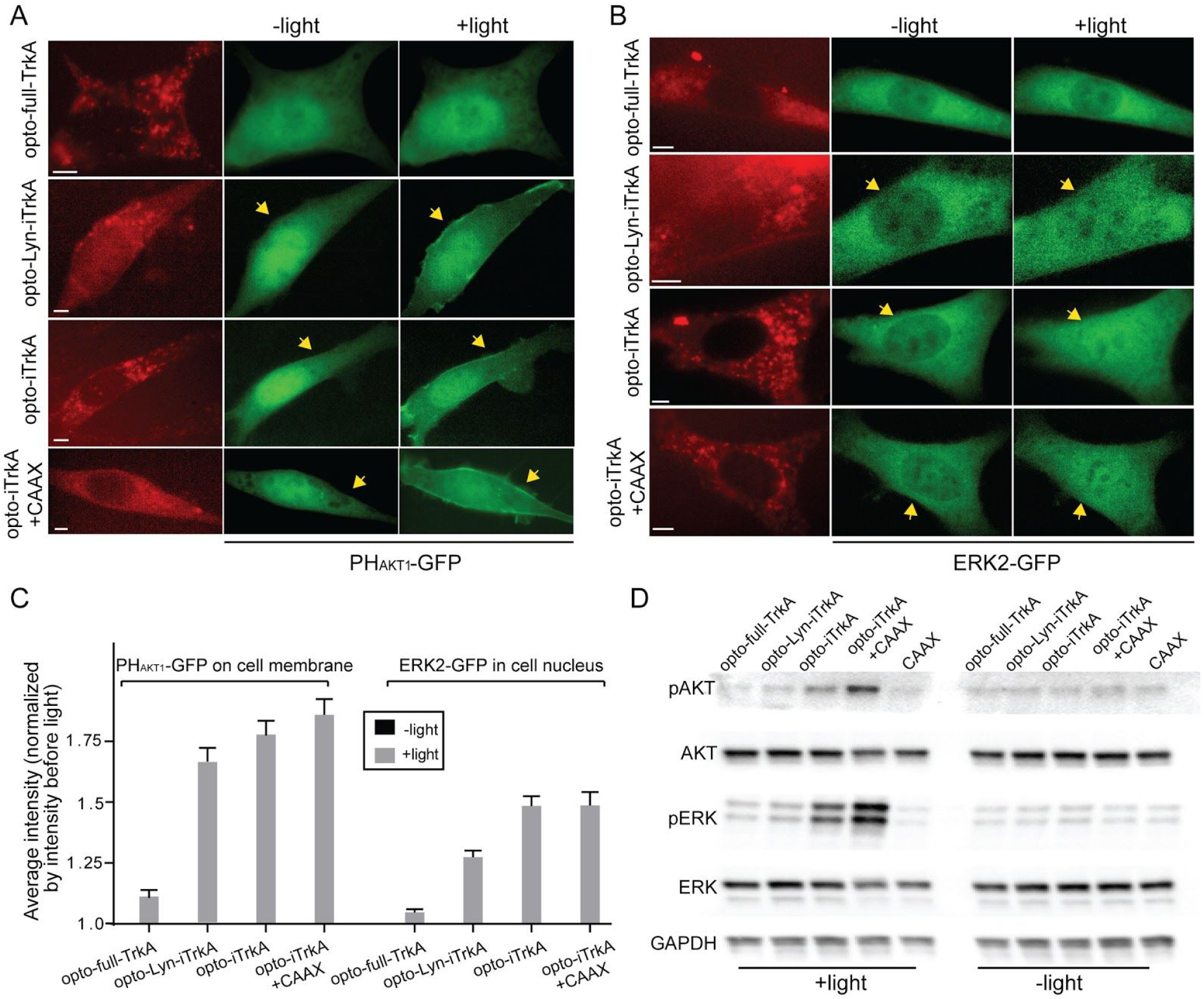
CRY2-TrkA systems activate PI3K/AKT and Raf/MEK/ERK signaling pathways. (A) Opto-Lyn-iTrkA, opto-iTrkA, and opto-iTrkA+CAAX exhibit activation of the PI3K/AKT signaling pathway, assayed by the membrane translocation of PH_AKT_-GFP in response to blue light stimulation. Opto-full-TrkA does not induce obvious PH_AKT_-GFP membrane translocation. (B) opto-Lyn-iTrkA, opto-iTrkA, and opto-iTrkA+CAAX systems induce ERK activation upon blue light stimulation, assayed by the nuclear translocation of ERK-GFP. Opto-full-TrkA does not induce obvious nuclear translocation of ERK-GFP. (C) Average translocation of PH_AKT_-GFP and ERK-GFP upon blue light stimulation for the four CRY2-TrkA systems. (for PH_AKT_-GFP assay, n=10,11,11,10 cells and for ERK-GFP assay, n=11,10,11,10 cells, respectively). (D) Western blot analysis of CRY2-TrkA systems for phosphorylated ERK (Thr202 and Tyr204) and phosphorylated AKT (Ser473). Opto-iTrkA and opto-iTrkA+CAAX induce AKT and ERK phosphorylation upon blue light stimulation. Scale bars, 5µm.

We further probed the light-induced activation of PI3K/AKT and Raf/MEK/ERK by immunoblot analysis of phosphorylated ERK (pERK1/2, Thr202 and Tyr204) and phosphorylated AKT (pAKT, Ser473). PC12 cells were transfected with each optoTrkA system and subjected to continuous blue light stimulation (200 µw/cm^2^) for 10 min. As a control, one group of cells was transfected with CIB1-GFP-CAAX alone. For an additional control, a second set of samples was subjected to the same transfection and analysis but kept in dark. Following light illumination, cells were immediately lysed and probed for pAKT, total AKT, pERK1/2, total ERK1/2, and GAPDH as a loading control. In the absence of blue light, very little phosphorylation of ERK or AKT was detected in any of the five conditions (Figure 2D). After blue light illumination, pAKT and pERK levels remain low in the CIB1-GFP-CAAX control. Cells expressing opto-Lyn-iTrkA, showed a slight increase in the levels of pAKT and pERK after blue light stimulation compared to the CAAX control. Blue light induced a significant increase in the levels of pAKT and pERK in cells expressing cytosolic opto-iTrkA, and an even greater increase in cells expressing opto-iTrkA+CAAX. In summary, our results demonstrate that three of our four optoTrkA strategies are capable of activating downstream Raf/ERK and PI3K/AKT signaling pathways, but that the extent of signaling is highly sensitive to the strategy used for activation. Specifically, blue light induced strong and robust activation of the downstream signaling in the opto-iTrkA or opto-iTrkA+CAAX systems, but much weaker activation in the opto-flTrkA and opto-Lyn-iTrkA systems.

We next examined whether light-gated activation of TrkA could simulate NGF in promoting neurite growth in PC12 cells. We used a subclone of the PC12 line, NeuroScreen-1 (NS1), which is widely used to assess neurite outgrowth in response to NGF treatment.^35^ Though our TrkA and iTrkA constructs include the fluorescent reporter tag mCherry, this fluorescence was often too dim to be used to visualize whole-cell morphologies under low magnification (20x), especially in fine neurite structures. Thus we made use of GFP to visualize the cell morphology. In the opto-iTrkA+CAAX system, this was achieved by using the fluorescently tagged CIB1-GFP-CAAX, and in all other systems by co-transfection with cytosolic GFP. PC12 cells transfected with CIB1-GFP-CAAX alone served as a control.

With blue light stimulation, PC12 cells expressing either the CIB1-GFP-CAAX control or opto-flTrkA failed to grow neurites (Figure 3A, E). On the other hand, cells transfected with opto-Lyn-iTrkA, opto-iTrkA or opto-iTrkA+CAAX grew out long neurites under blue light stimulation (Figure 3B-D). Among them, cells expressing opto-iTrkA+CAAX grew out significantly longer neurites than other groups. The transfected cells kept in dark did not grow measurable neurites, except for a few cells expressing opto-Lyn-iTrkA that showed some neurite outgrowth in dark (Figure S3). To quantify the extent of PC12 cell differentiation, the percentage of cells with neurites were measured among all the transfected cells either in dark or under light illumination (Figure 3F, statistics shown in Table S1). Under blue light, 34% of opto-iTrkA and 61% of opto-iTrkA+CAAX cells grew neurites longer than the size of their cell bodies, significantly more than their dark controls (6.2% and 1.7%, respectively). 22% of opto-Lyn-iTrkA cells grew neurites with light stimulation, but this growth is not significantly greater than the corresponding dark control (13%).

**Figure 3:**
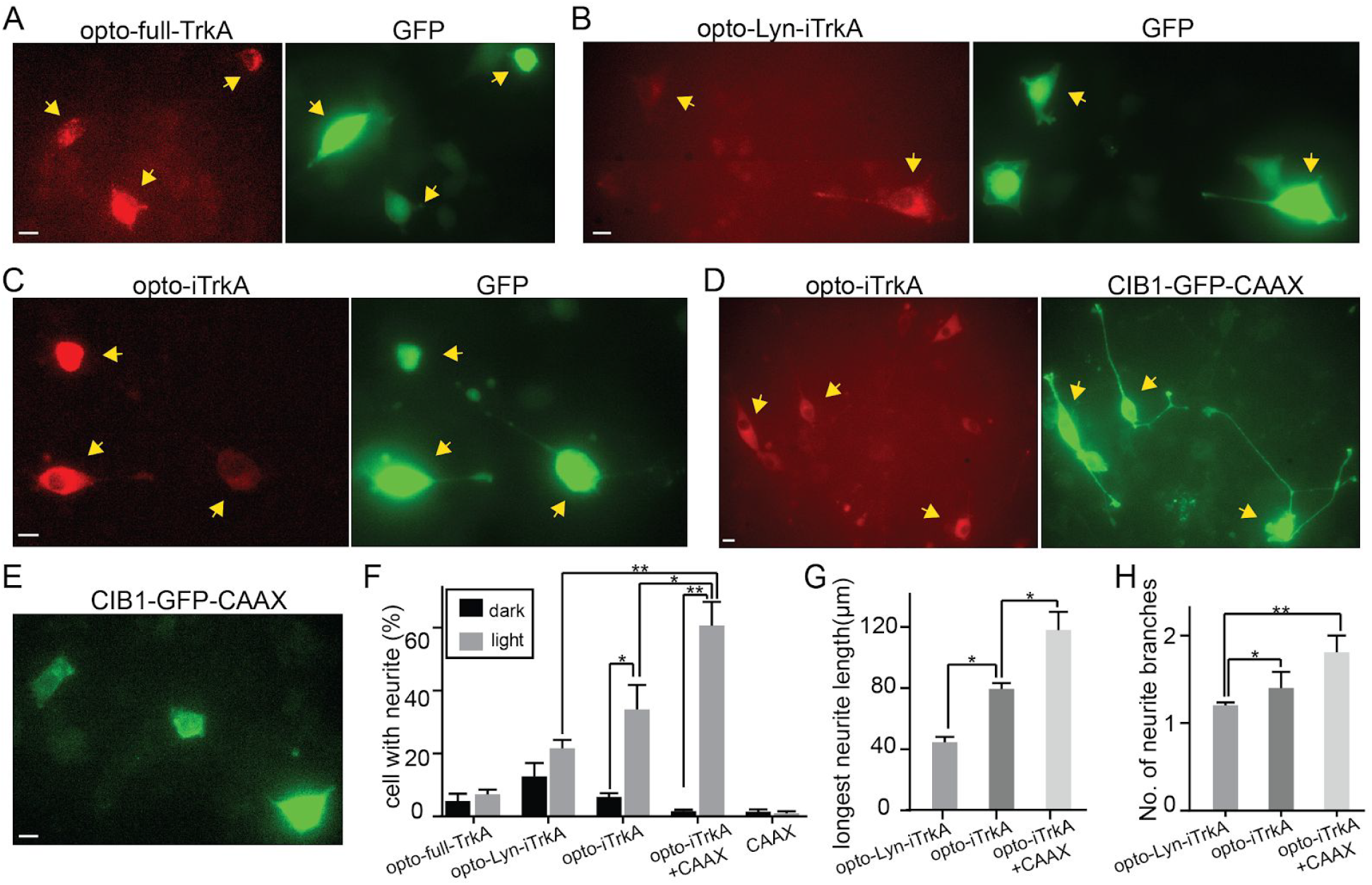
Light-induced activation of TrkA promotes neurite growth in PC12 cells in the absence of NGF. PC12 cells were subjected to blue light illumination at 200 mW/cm^2^ for 24hr. (A) PC12 cells expressing opto-full-TrkA and GFP displayed no obvious neurite growth. (B) Some PC12 cells expressing opto-Lyn-iTrkA and GFP extend short neurites. (C) Some PC12 cells expressing opto-iTrkA and GFP exhibit neurite growth. (D) Most PC12 cells expressing opto-iTrkA+CAAX grow long neurites. (E) PC12 expressing only CIBN-GFP-CAAX did not show noticeable neurite growth. (F) Quantification of percentage of cells with neurite growth for different systems. (G) Quantification of the length of the longest neurite in cells bearing neurites. (H) Quantification of the average number of neurites in cells bearing neurites. Results are averaged from three independent sets of experiments (detailed in **Supplementary Table 1**) and presented as mean ± SEM. Statistical analysis was performed using one-way ANOVA with Dunnett’s post hoc test. Student’s T-test was used between dark or light cells under the same transfection conditions. (*P<0.05, **P<0.005) Scale bars, 10µm.

We also measured the average length of the longest neurite of each cell, which was greatest in cells expressing opto-iTrkA+CAAX (118 µm) and shorter in cells transfected with opto-iTrkA (80 µm) and cells transfected with opto-Lyn-iTrkA (44 µm) (Figure 3G). Finally, the average numbers of neurites per cell were quantified to compare the morphological differences (Figure 3H). Cells expressing opto-iTrkA+CAAX had more neurite branches than other conditions. In summary, these results showed that light-induced activation of opto-iTrkA+CAAX and opto-iTrkA is sufficient to induce neurite growth in PC12 cells. In particular, opto-iTrkA+CAAX activation is the most effective in promoting neurite growth in PC12 cells, and exhibits neurites of higher complexity than our other systems.

As NGF promotes neuronal survival through TrkA, we next tested whether light-activated TrkA signaling could mimic NGF function in supporting neuronal survival. We used embryonic DRG neuron culture as a model system, as a large proportion of these cells, known as peptidergic nociceptors, are not viable in the absence of NGF.^36–38^ DRG neurons were transfected with each optoTrkA system by electroporation at day 0. We used the human synapsin 1 promoter^39^ to express GFP or CIB1-GFP-CAAX exclusively in neurons in order to avoid high glial background expression. After transfection, neurons were plated in a 12-well plate and allowed to recover for 16 hours in complete medium supplemented with 50ng/ml NGF. NGF was then withdrawn from the culture medium and cells were treated with anti-NGF antibody to remove residual growth factor for 48 hr. During this withdrawal, the cultures were subjected to blue light illumination at 200 uW/cm^2^ using home-made LED arrays. Another group of transfected neurons were kept in dark as controls. After 48 hr in the presence or absence of blue light stimulation, the cultures were fixed and immunostained against GFP to visualize the neuron morphologies.

DRG neurons expressing opto-flTrkA did not survive either under light illumination or in dark (Figure 4A). A small percentage of DRG neurons transfected with opto-Lyn-iTrkA (28%) survived in the presence of blue light stimulation (Figure 4B). Surviving DRG neurons had extended out long axons. On the other hand, a substantial number of DRG cells expressing opto-iTrkA (57%) or opto-iTrA+CAAX (73%) survived after light illumination and grew very long neurites (Figure 4C and 4D). For control samples kept in dark, very few cells survived in each group of transfected DRG. To quantify the efficacy in promoting neuronal survival, the survival rates were measured for each transfection in dark or under light stimulation (Figure 4E, Table S2). Transfected DRG neurons with long neurites were identified as living cells. Quantitative analysis confirmed that opto-iTrA+CAAX is the most effective in supporting DRG survival in the absence of NGF, followed by opto-iTrkA and opto-Lyn-iTrkA. Opto-full-TrkA did not support DRG survival. This neuron survival result corroborates the neurite growth studies in PC12 cells to show that light-induced activation of opto-iTrkA+CAAX is highly effective in mimicking endogenous NGF functions.

**Figure 4:**
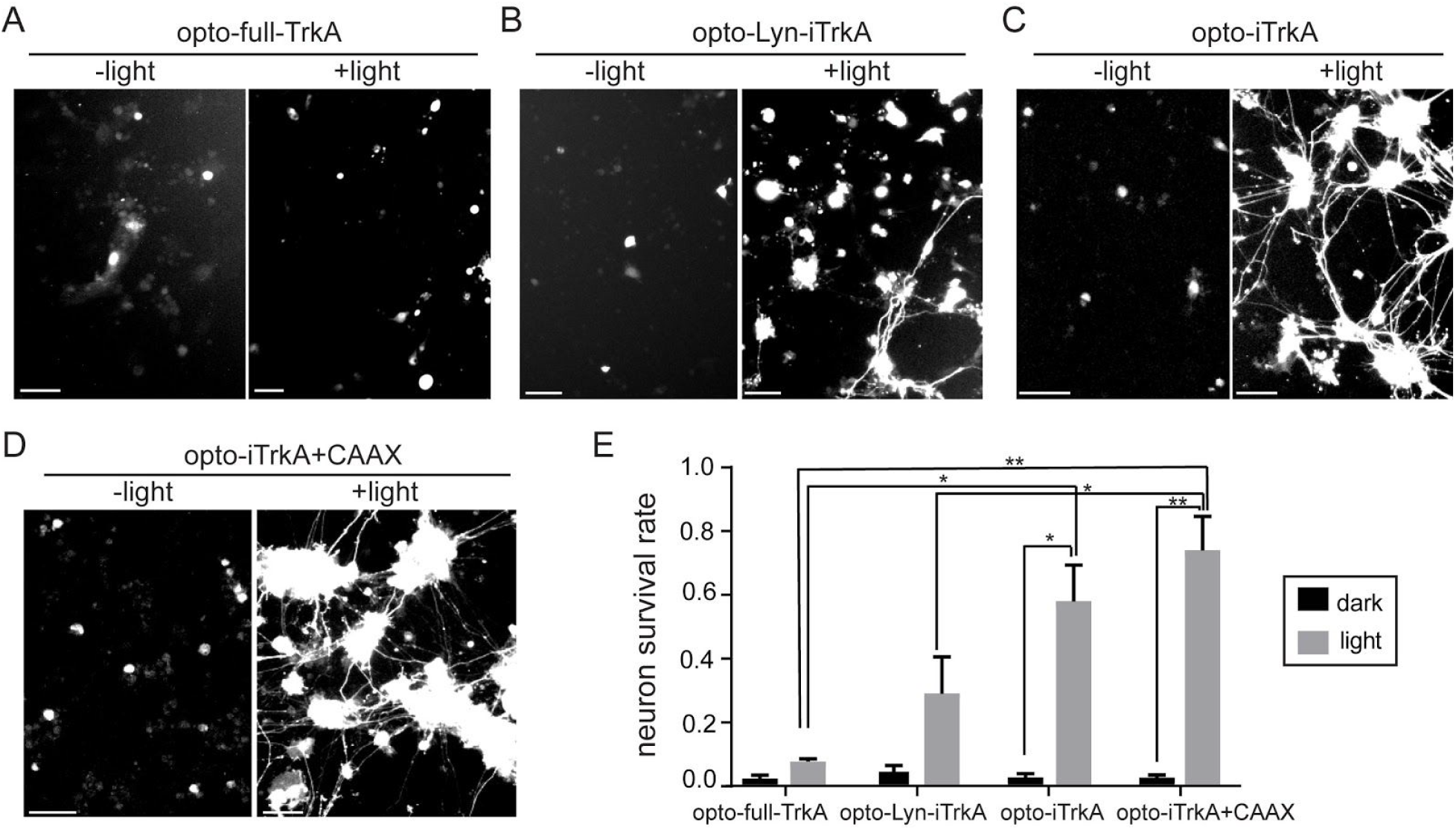
Light-induced TrkA activation supports DRG neuron survival in the absence of NGF. After transfection, primary rat DRG neurons were allowed to recover in NGF medium for 24hrs. Then, NGF was withdraw and replaced with anti-NGF for 48hrs while the cultures were subjected to blue light stimulation or kept in dark for 2 days. (A) DRG neurons transfected with opto-full-TrkA and GFP did not survive under light or dark. (B) Some DRG neurons expressing opto-Lyn-iTrkA and GFP survived upon blue light stimulation, but few survived in the dark control. (C) Many DRG cells expressing opto-iTrkA survived and had extended long axons upon blue light stimulation, but not in the dark controls. (D) Most DRG neurons expressing opto-iTrkA and CIB1-GFP-CAAX were alive with long axons with blue light, but not in the dark controls. (E) quantification of survival rate of DRG neurons for TrkA/CRY2 systems, in light and dark conditions. Results are averaged from three independent sets of experiments (see numbers of cells in each set of experiment in **Table S2**) and presented as means ± SEM. One-way ANOVA with Dunnett’s post hoc test was performed between cells with different transfection. T-test was performed between cells in dark or in light with the same transfection. (*P<0.05, **P<0.005) Scale bars, 50µm.

In this work, we designed and evaluated four strategies to achieve light-induced TrkA signaling either in the cytosol or on the cell membrane. Cytosolic activation is achieved by fusing CRY2 to intracellular domain of TrkA (opto-iTrkA) so that the light-mediated homo-oligomerization of CRY2 can result in the oligomerization and autophosphorylation of iTrkA. To enable light-inducible TrkA activation at the plasma membrane, we tested three strategies: opto-flTrkA, opto-Lyn-iTrkA, and opto-iTrkA+CAAX. We find that opto-iTrkA+CAAX is the most effective in activating downstream TrkA signaling pathways, stimulating neurite outgrowth in PC12 cells, and supporting DRG survival in the absence of NGF.

Chang *et al.*’s work ^26^ demonstrated the feasibility of using CRY2-based optogenetic protein dimerization to build light-inducible neurotrophin receptors. Here, we sought to validate an optoTrkA system that could reproduce more complex signaling outcomes, such as differentiation and survival. We have also demonstrated that strategies which make use of only the intracellular domain of TrkA are more effective at recapitulating native signaling than strategies that utilize the full length receptor. One possible explanation for this difference is that full length TrkA may exist as pre-formed yet inactive dimers prior to ligand binding.^40^ The extracellular domain and transmembrane domain of full-TrkA may lock the pre-formed dimer in a configuration that cannot be fully activated by CRY2 oligomerization. Further, the separation of the intracellular, signaling-competent domain of TrkA from its ligand-binding extracellular domain confers its own advantage, especially for use in model systems which may produce NGF endogenously. By removing the ligand-binding extracellular domain, we render the optogenetic construct inert in the absence of light induction, thus increasing confidence that any observed change is due to our perturbation rather than background.

Among the iTrkA systems, we observed that opto-Lyn-iTrkA is less efficient in inducing cell differentiation and supporting neuronal survival than opto-iTrkA+CAAX. opto-Lyn-iTrkA also showed higher background activities in dark than other systems, e.g. cells transfected with opto-Lyn-iTrkA had some neurite growth in dark, likely due to the enhanced homo-oligomerization of membrane-bound CRY2, which has been reported in a previous study.^41^

In summary, we have successfully constructed and validated two optoTrkA systems, opto-iTrkA+CAAX and opto-iTrkA, to optimally induce downstream TrkA signaling pathways. In particular, opto-iTrkA+CAAX is the most effective in mimicking the role of NGF in stimulating neurite growth in PC12 cells and supporting DRG neuronal survival. The opto-iTrkA+CAAX strategy allows precise spatial and temporal control of TrkA signaling to facilitate the study of mechanisms underlying NGF/TrkA signaling in a quantitative manner. In addition, the opto-iTrkA+CAAX strategy might be a useful tool to screen therapeutics towards diseases where NGF/TrkA signaling is critically involved, especially in instances which require high spatiotemporal precision.

## Methods

### Plasmid construction

TrkA-mCh-CRY2 (opto-full-TrkA): Full length TrkA was inserted into pmCherry-N1 (clontech) at XhoI and AgeI sites. TrkA template plasmids were described previously [22]. CRY2 was inserted into pTrkA-mCherry with a GS N-terminal linker at BsrG1 and MfeI using InFusion (Clontech).

CRY2-mCh-iTrkA (opto-iTrkA): The intracellular domain of TrkA (iTrkA, aa 450-799, NP_067600.1) with GSS linker was inserted into CRY2-mCh at BsrG1 and MfeI using InFusion (Clontech).

Lyn-iTrkA-mCh-CRY2 (opto-Lyn-iTrkA): Lyn sequence (MGCIKSKRKDNLNDD) was directly synthesized (Integrated DNA Technology) and inserted into pmCherry-N1 at NheI and XhoI sites using ligation method (Thermo Fisher). iTrkA was then inserted at XhoI and AgeI and CRY2 was inserted at BsrG1 and MfeI using InFusion (Clontech).

hSyn-CIB1-GFP-CAAX and hSyn-GFP: human synapsin promoter (hSyn) sequence was obtained by PCR from vector pLV-hSyn-RFP (Addgene #22909) and inserted into CIB1-GFP-CAAX [7] or pEGFP-N1 (Clontech) at VspI and NheI using InFusion (Clontech).

### Cell culture and transfection

PC12 cells (NeuroScreen-1 subclone) were maintained in complete medium (F12K supplemented with 15% horse serum and 2.5% FBS). NIH 3T3 cells were grown in DMEM medium supplemented with 10% fetal bovine serum (FBS). All cell cultures were maintained a standard incubator at 37°C with 5% CO_2_. Both PC12 and 3T3 cells were transfected with desired DNA plasmids using Lipofectamine 2000 (Life Technologies) according to the manufacturer’s protocol. The transfected cells were allowed to recover and express the desired proteins overnight in culture medium. PC12 cells were kept in starvation medium (F12K only) overnight before light stimulation. For ERK and AKT translocation assays, 3T3 cells were starved in DMEM only for overnight before imaging.

DRG neurons were dissected from E18 embryos by standard procedure [23] and maintained on poly-L-lysine (Sigma) coated coverslips (VWR) in DRG maintenance medium (Neurobasal supplemented with GlutaMax and B27, and containing 50 ng/ml NGF). For NGF withdrawal, 50ng/ml NGF was replaced with 100ng/ml NGF antibody (Millipore AB1526SP). The Amaxa™ Nucleofector™ was used for electroporation as previously described.^42^

### Programmable LED light box

LED light boxes were constructed as previously reported for long-term light illumination.^43^ Briefly, a 12 well plate sized blue LED array was constructed by assembling 12 blue LEDs (LED465E, ThorLabs) on a breadboard, housed in an aluminum box, and a light diffuser film was positioned above the LED array to ensure uniform light intensity in defined area, separated by black barriers. The light intensity of each LED was controlled through a tunable resistor, and was measured by a power meter (Newport, 1931-C). The LED array was controlled by Labview through a data acquisition board (National Instrument-DAQ, PCI-6035E). The LEDs were supplied with user-defined DC voltages that were individually controllable through the Labview program.

### Optical activation in culture

For PC12 neurite growth experiments, transfected PC12 cells were kept in dark and serum-starved for 24 hours prior to blue light stimulation to minimize interference from growth factors present in serum. Cells were then illuminated with continuous blue light at 200 µw/cm^2^ for 24 hours using a custom-built LED array housed inside a CO_2_ incubator. This illumination condition has been repeatedly tested and shows no obvious toxicity as demonstrated in our previous studies.^23,41,43^ Additional sets of transfected cells were kept in dark as controls. After 24 hours, cultures were fixed and mCherry and GFP were visualized using fluorescence microscopy.

For DRG survival experiments, transfected neurons were plated in a 12-well plate and allowed to recover for 16 hours in complete medium supplemented with 50ng/ml NGF. NGF was then withdrawn from the culture medium and cells were treated with anti-NGF antibody to remove residual growth factor for 48 hr. During this withdrawal, the cultures were subjected to blue light illumination at 200 uW/cm^2^ using home-made LED arrays. Another group of transfected neurons were kept in dark as controls. After 48 hr in the presence or absence of blue light stimulation, the cultures were fixed and immunostained against GFP to visualize the neuron morphologies.

### Immunoblotting

Immunoblotting experiments were carried out as previously described.^44^ Briefly, after desired treatment, cultured cells were lysed in PB/Triton buffer (7.74 mM Na_2_HPO_4_, 2.26 mM NaH_2_PO_4_, 5 mM EDTA, 5 mM EGTA, 100 mM NaCl, 10 mM Na_4_P_2_O_7_, 50 mM NaF, 1 mM Na_3_VO_4,_ 1% Triton X-100), and clarified lysates were mixed with Laemmli sample buffer (Bio-Rad 1610747) and β-mercaptoethanol and boiled for 10 minutes. Lysed samples were subjected to electrophoresis using Bio-Rad’s Mini-PROTEAN system (1658026FC). After separation, protein was transferred to a polyvinylidene fluoride membrane (Bio-Rad 1704273), followed by standard blotting procedure. Primary antibodies were obtained from Cell Signaling Technology: anti-pAkt (T308) (CST 9275S), anti-Akt (CST 9272), anti-pErk1/2 (T202+Y204) (CST 9101S), anti-Erk1/2 antibody (CST 9102S), and anti-GAPDH (CST 2118). HRP-conjugated secondary antibody (CST 7074) was used for protein band detection. Protein bands were visualized by chemiluminescence (Bio Rad 1705060).

### Cell fixation and immunocytochemistry staining

For PC12 neurite growth assays and DRG survival assays, cells were washed with ice-cold PBS three times before fixation in 4% paraformaldehyde in sucrose-buffered solution at room temperature for 20 minutes. The fixed cells were then washed with ice-cold PBS twice and proceeded to imaging or immunostaining.

DRG neurons were stained with anti-GFP (MolecularProbe, #A11122) in 5% BSA-PBS buffer at 4°C overnight, and then probed using Alexa Fluor 488 (Thermo Fisher, R37116).

### Fluorescence Imaging

Live cell imaging was performed on an epifluorescence microscope (Leica DMI6000B) equipped with an on-stage CO_2_ incubation chamber (Tokai Hit GM-8000) and a motorized stage (Prior). An adaptive focus control was used to actively keep the image in focus during the period of imaging. A light-emitting diode (LED) light source (Lumencor Sola) was used as the fluorescence light source. For blue-light stimulation, pulsed blue light (200 ms pulse duration at 9.7 W/cm^2^) was used for GFP imaging and to initiate CRY2-CIB1 and CRY2-CRY2 interactions. For mCherry imaging, pulsed green light (200 ms pulse duration) was used. Fluorescence signal from GFP was detected using a commercial GFP filter cube (Leica; excitation filter 472/30, dichroic mirror 495, emission filter 520/35); fluorescence signal from mCherry was detected using a commercial Texas red filter cube (Leica; excitation filter 560/40, dichroic mirror 595, emission filter 645/75). The images for ERK or AKT translocation were acquired with an oil-immersion 100× objective while the images for PC12 or DRG morphologies were acquired with 10× or 40× objectives. All experiments were imaged with a sensitive CMOS camera (PCO.EDGE 5.5) (PCO).

### Data analysis

To analyze the neurite growth of PC12 cells, neurites were measured manually using the ImageJ line tool, and neurites significantly shorter than cell body were not considered. The number of cells in each measurement are included in Supplementary Table 1.

To quantitatively analyze the translocation of PH_AKT1_ to plasma membrane, average intensity of membrane I_membrane_ was measured by selecting and measuring only membrane area. Average intensity of cytosol intensity I_cytosol_ was measured by selecting the whole cytosol area. I_background_ is defined as the average intensity of an image from blank areas (outside cells). The amount of PH_AKT1_ on membrane was indicated by ratio=(I_membrane_ – I_background_)/(I_cytosol_ – I_background_). In Figure 2C, the normalized average intensity of PH_AKT1_-GFP was calculated by ratio_+light_/ratio_-light_.

Similarly, the average intensity of ERK2-GFP in the nucleus was quantified by ratio=(I_nucleus_ – I_background_)/(I_cytosol_ – I_background_). In Figure 2C, the normalized average intensity of ERK2-GFP was calculated by ratio_+light_/ratio_-light_.

## Author Contribution

BC, SG, and LD conceived the project and designed the experiments. SG, LD, and JMH performed experiments, and LD analyzed data. BC, SG, LD, and JMH wrote the manuscript and all authors edited the manuscript.

## Acknowledgement

We thank Dr. Chandra Tucker (University of Colorado, Denver) for providing CIB1-GFP-Caax and CRY2-mCherry and Dr. Tobias Meyer (Stanford University) for providing PH_AKT1_-GFP plasmid. We also thank Zhuoluo Feng (Stanford University) for his help in constructing the controllable blue LED array. GS is a New York Stem Cell Foundation Robertson Investigator. This work was supported by the US NIH (DP2-NS082125), a Packard fellowship in Science and Engineering, a National Science Foundation Graduate Research Fellowship under Grant No. DGE-1656518 (JMH), and a seed grant from the Stanford Neurosciences Institute (BC and GS).

## Supporting Information Available

Figures S1-S3 and Tables S1-S2. This material is available free of charge via the Internet at http://pubs.acs.org.

