## Supplementary Materials for "Optical activation of TrkA signaling"

**Figure S1**

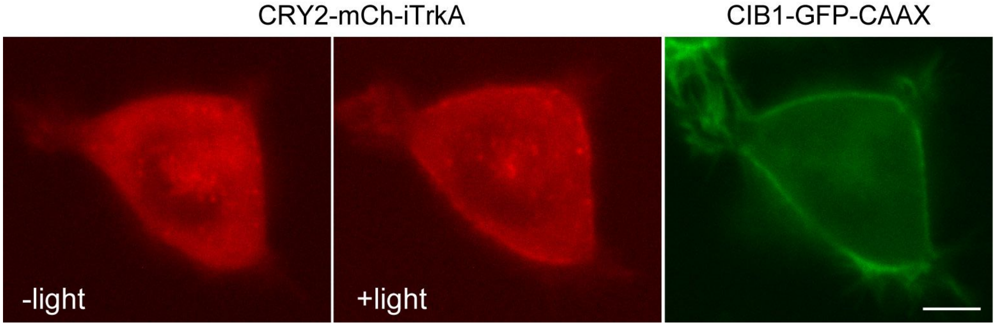


Figure S1: CRY2-mCh-iTrkA (opto-iTrkA) can be succesfully recruited to plasma membrane by binding with CIB1-GFP-CAAX. PC12 cells were co-transfected with CRY2-mCh-iTrkA and CIB1-GFP-CAAX and subject to one pulse of 200 ms blue light exposure. Scale bar, 5µm.

**Figure S2**

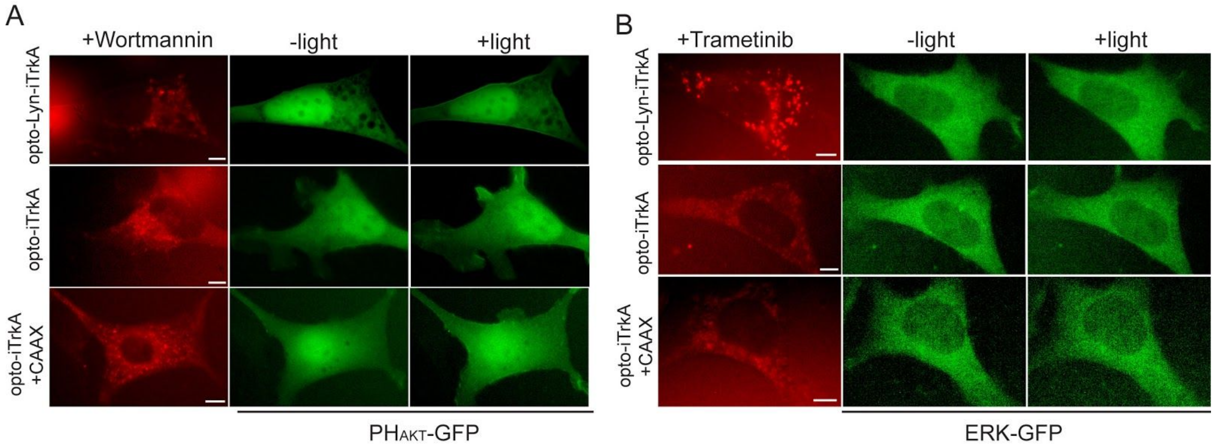


Figure S2: PI3K inhibitor wortmannin or MEK inhibitor trametinib suppressed the translocation of $PH_{AKT}$-GFP or ERK-GFP in cells expressing TrkA/CRY2 systems. 3T3 cells were transfected with opto-Lyn-iTrkA, opto-iTrA or opto-iTrA+CAAX as well as $PH_{AKT}$-GFP (A) or ERK-GFP (B). In cells expressing opto-iTrkA+CAAX, CIB1-CAAX was used without fluorescence reporter. The redistribution of $PH_{AKT}$-GFP or ERK-GFP was suppressed compared with cells without the addition of inhibitors. Scale bars, 5µm.

**Figure S3**

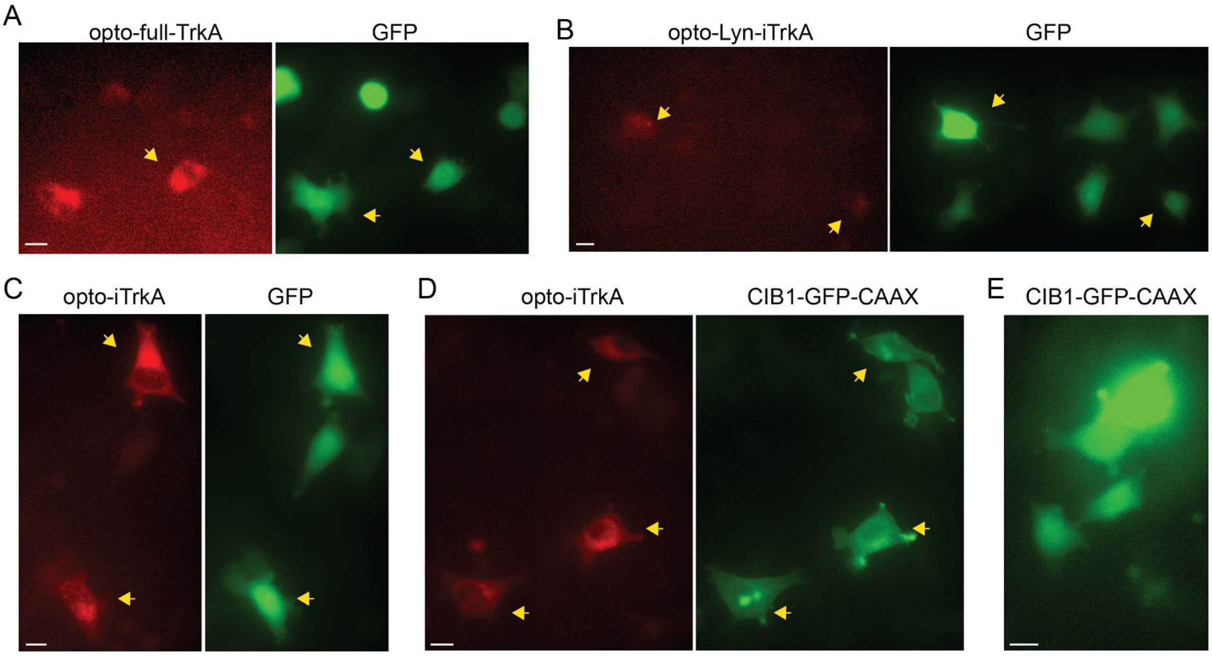


Figure S3: In dark, PC12 cells expressing each of the four TrkA/CRY2 systems(A-D) or CIB1-GFP-CAAX (E) showed no obvious neurite growth. Scale bars, 10μm.

**Table S1**

| | | opto-full-TrkA | | opto-Lyn-iTrkA | | opto-iTrkA | | opto-iTrkA+CAX | | CAAX only | |
|---|---|---|---|---|---|---|---|---|---|---|---|
| | #of cells | Total # | + neurite | Total # | + neurite | Total # | + neurite | Total # | + neurite | Total # | + neurite |
| Set 1 | dark | 286 | 4 | 290 | 14 | 306 | 12 | 289 | 2 | 278 | 6 |
| | light | 243 | 9 | 199 | 53 | 292 | 140 | 194 | 142 | 293 | 6 |
| Set 2 | dark | 262 | 13 | 300 | 40 | 306 | 21 | 289 | 8 | 255 | 1 |
| | light | 272 | 22 | 285 | 55 | 238 | 52 | 181 | 113 | 265 | 3 |
| Set 3 | dark | 312 | 28 | 301 | 60 | 341 | 27 | 289 | 4 | 261 | 7 |
| | light | 231 | 21 | 275 | 53 | 230 | 76 | 208 | 98 | 292 | 2 |

Table S1: number of PC12 cells for quantification in Fig. 3

**Table S2**

| | | opto-full-TrkA | | opto-Lyn-iTrkA | | opto-iTrkA | | opto-iTrkA+CAX | |
|---|---|---|---|---|---|---|---|---|---|
| | #of cells | Total # | surive d | Total # | surive d | Total # | surived | Total # | surived |
| Set 1 | dark | 122 | 1 | 179 | 2 | 96 | 0 | 194 | 8 |
| | light | 366 | 33 | 197 | 79 | 422 | 333 | 701 | 615 |
| Set 2 | dark | 119 | 5 | 94 | 8 | 194 | 8 | 99 | 1 |
| | light | 76 | 6 | 223 | 11 | 88 | 47 | 226 | 119 |
| Set 3 | dark | 136 | 1 | 75 | 4 | 132 | 3 | 70 | 1 |
| | light | 77 | 5 | 103 | 41 | 145 | 58 | 120 | 96 |

Table S2: number of DRG cells for quantification in Fig. 4
